# A single-plane electrical impedance tomography framework for integrated liver and kidney assessment

**DOI:** 10.64898/2026.10.26.762009

**Authors:** Aywin Wu, Jen-Chien Lai, Mo Kwan Ho, Justin Q.J. Ge, Nan-jie Gong, Iris Y. Zhou, Monica Wong, Bradley J. Edelman, Russell W. Chan

## Abstract

Chronic liver and kidney diseases frequently coexist, but their assessment typically requires separate clinical workflows involving serum-based estimated glomerular filtration rate (eGFR) testing and liver elastography-based controlled attenuation parameter (CAP). Electrical impedance tomography (EIT) has recently shown promise for providing portable, non-invasive proxies for both reference measurements. However, existing multi-organ EIT assessment still requires different acquisition protocols for each organ. To simplify this procedure, we assessed the feasibility of a single-plane EIT framework for reconstructing organ-specific images and estimating liver and kidney health from one abdominal belt placement. Across diverse abdominal belt locations, opposite-plane deviation was significantly greater for eGFR_EIT_ estimates (2.92%; IQR, 0.46-12.53%) than for CAP_EIT_ estimates (0.94%; IQR, 0.66-1.70%) (p < 0.01). For both organs, compared to images acquired at the conventional organ-specific planes, image correlation significantly decreased (both, p < 0.05) and normalized root mean squared error significantly increased (both, p < 0.05) at the furthest imaging plane, with these differences being numerically larger for the kidney. Together, these results supported the kidney plane as the preferred location for multi-organ assessment. Test-retest evaluation after belt removal and reapplication demonstrated that the correlation between two repeated CAP_EIT_ estimates was just as strong at the kidney plane (R^2^ = 0.98, p < 0.001) as it was at the conventional liver plane (R^2^ = 0.97, p < 0.001). Finally, in an independent clinical cohort, kidney-plane CAP_EIT_ estimates were significantly higher in participants with a high metabolic liver-risk profile (i.e. diabetes and high blood pressure) (p < 0.005). CAP_EIT_ estimates also distinguished high and low metabolic risk groups with an AUC of 0.756. Together, these findings support the feasibility of a portable multi-organ EIT imaging framework capable of extracting complementary liver and kidney information from a single belt placement.

## Introduction

Chronic liver and kidney diseases frequently develop in the same metabolic context, yet both may remain silent until clinically meaningful dysfunction has developed. Metabolic dysfunction-associated steatotic liver disease (MASLD) is now among the most prevalent chronic liver conditions worldwide, affecting approximately one-third of adults ^1^. Chronic kidney disease (CKD) similarly affects a substantial and growing fraction of the adult population ^2,3^. Beyond sharing risk factors such as diabetes, hypertension, obesity, and cardiovascular disease, MASLD and CKD frequently coexist and may contribute to dual-disease progression ^4–6^. This convergence provides a rationale for assessing liver and kidney risk together rather than through serial single-organ evaluations. In fact, population imaging and routine diagnostic studies increasingly demonstrate the feasibility of combining measurements from multiple organs for integrated risk assessment ^7,8^. Approaches capable of collecting complementary liver and kidney information during one encounter could therefore support primary-care risk assessment, preventive care, and longitudinal health monitoring while reducing reliance on repeated specialist examinations.

Current medical assessment pathways provide valuable but often fragmented information. A single laboratory encounter can generate biochemical markers relevant to both the liver and kidneys, but routine in-vitro testing does not directly quantify hepatic steatosis or provide imaging-resolved kidney information ^9,10^. In addition, although imaging modalities such as ultrasound and MRI can examine multiple abdominal organs, common quantitative endpoints utilized in the clinic (e.g. fat content, fibrosis) are generally obtained through very dissimilar means with varying accuracy levels ^11,12^. These inconsistencies are highlighted in the context of gold standard clinical assessments of the liver and kidney where hepatic steatosis involves quantification using magnetic resonance proton-density fat fraction (MR-PDFF) or vibration-controlled transient elastography (VCTE)-based controlled attenuation parameter (CAP) ^13–15^, and kidney function is evaluated using serum-based estimates of glomerular filtration rate (eGFR) and markers of kidney damage using the albumin-to-creatinine ratio (uACR) ^10,16^. Comprehensive multi-organ assessment may therefore span laboratory collection, imaging or elastography services, specialist interpretation, and subsequent clinical follow-up. Access to advanced imaging and liver elastography also remains uneven, particularly outside traditional hospital settings and in resource-constrained areas ^17,18^. A portable imaging platform that is capable of acquiring organ-specific information could complement rather than replace confirmatory testing by identifying individuals who warrant further assessment for individual or multiple organ systems.

Electrical impedance tomography (EIT) is a non-invasive imaging modality that estimates internal conductivity distributions from boundary voltage measurements obtained during the application of small alternating currents ^19^. The technique is portable, non-invasive, and radiation free, and is capable of repeated measurements at the point of care by non-specialized personnel ^20^. The most mature clinical application of EIT is regional lung monitoring ^21–23^, however, recent advancements involving artificial intelligence have markedly expanded its application repertoire to multiple other organs in and outside of the clinic ^24^. Conventional time-difference EIT, as is typically used for continuous pulmonary monitoring, emphasizes conductivity changes over time that may help identify regional ventilation deficits. By contrast, frequency-difference EIT (fdEIT) exploits the frequency dependence of biological tissue impedance. Because tissue composition, extracellular and intracellular compartments, and cell membranes shape this spectral response, fdEIT can provide contrast that is relevant to abdominal tissue characterization ^25–27^. Importantly, prior studies have reported altered impedance signatures in both MASLD and CKD, which support further development of organ-specific fdEIT measurements ^28–30^.

To help expand access to organ health screening using such EIT features, we previously developed multi-frequency EIT analytics pipelines and machine learning models for estimating liver CAP (CAP_EIT_) and kidney eGFR (eGFR_EIT_) ^20,31–33^. These models combine quality-controlled EIT processing with basic anthropometric variables to return continuous estimates of the corresponding clinical measurements. In separate clinical cohorts (n > 200 patients total), these models achieved significant alignment with the reference values of CAP (R^2^ = 0.74) and eGFR (R^2^ = 0.57). Moreover, applying these models to detect the at-risk status of liver and kidney health achieved a sensitivity and specificity of 91.7% and 88.1% for liver risk, and 100% and 94% for kidney risk, respectively ^20,31,32^. Nevertheless, the original protocols associated with these works were validated with the electrode belt being placed at different craniocaudal levels for liver and kidney acquisition. For multi-organ imaging, repositioning the belt for each organ would increase acquisition time and introduce additional opportunities for placement error ^24,34^.

The anatomical proximity of the liver and kidneys raises the practical question of whether one abdominal belt position provides sufficient information for accurately estimating health metrics of both organs. In fact, EIT does not measure a pure two-dimensional imaging plane, but rather a lens-shaped region that includes conductivity changes from slightly above and below electrode belt location ^35^. Therefore, it is plausible that sufficient conductivity information from one organ is present in a nearby but anatomically distinct imaging plane. The primary goal of this study was to determine whether a common abdominal belt placement contains sufficient organ-specific information to support both liver and kidney assessment without belt repositioning.

Here, we tested a single-plane strategy through a sequence of complementary analyses (**Fig. 1**). We first quantified the sensitivity of liver and kidney health estimates to three adjacent belt locations, and then examined whether cross-plane fdEIT image concordance explained any organ-specific differences. We next assessed repeatability after belt removal and reapplication at the selected single plane, and finally, examined whether liver estimates from that plane retained clinically relevant separation between contrasting metabolic risk profiles.

**Figure 1.**
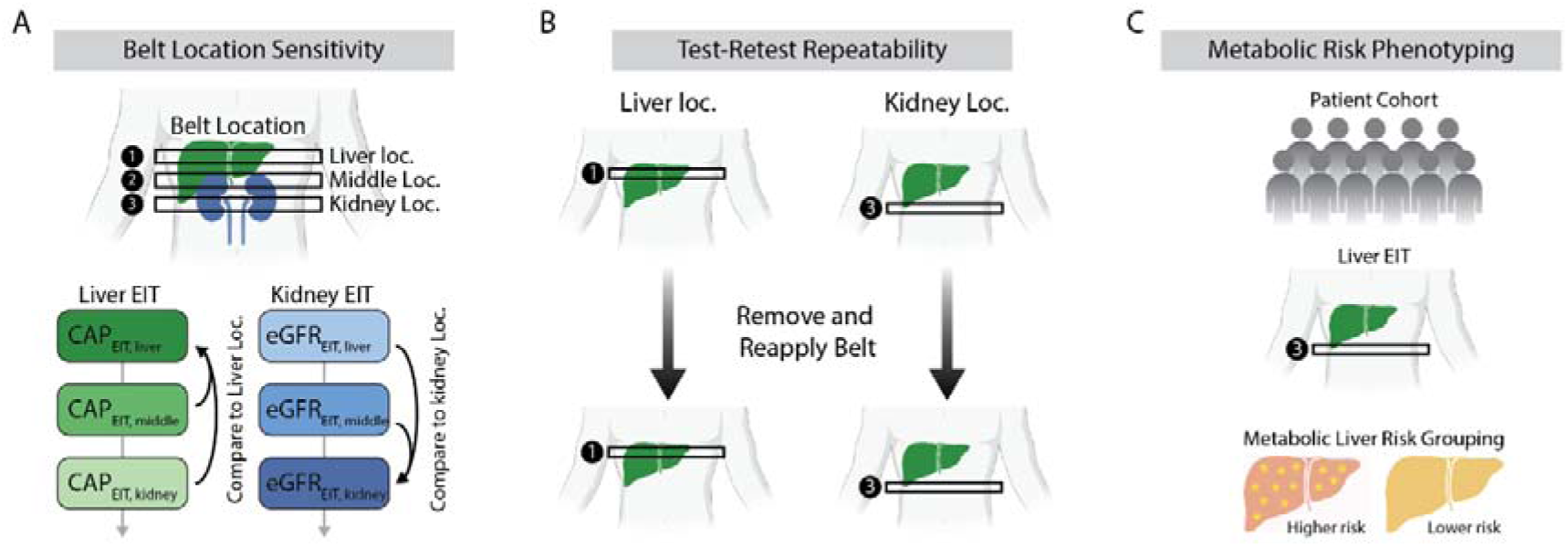
Study design for evaluating a common EIT belt plane for liver and kidney assessment. **(A)** Belt-location sensitivity was evaluated in 42 participants by acquiring liver and kidney EIT measurements at three adjacent craniocaudal positions. CAP_EIT_ estimates at the middle and kidney positions were compared with the conventional liver position (Pos 1), whereas eGFR_EIT_ estimates at the middle and liver positions were compared with the conventional kidney position (Pos 3). **(B)** Test-retest repeatability was evaluated in 15 participants who underwent repeated liver EIT measurements at the liver position and the kidney position, with complete belt removal and reapplication between acquisitions. **(C)** Clinical relevance was explored using EIT measurements from the kidney location in an independent cohort of 75 participants. Liver CAP_EIT_ estimates were compared between participants with contrasting metabolic liver-risk phenotypes.

## Results

### Belt location differentially affects liver and kidney health estimates

We first asked whether moving the belt away from each organ’s conventional acquisition plane changed its downstream health estimate. Here, we defined three imaging planes relative to the lowest rib: position 1 above the lowest rib (conventional liver plane), position 2 at the lowest rib, and position 3 at the inferior costal margin (conventional kidney plane). When examining valid liver and kidney health metrics across all three locations (**Fig. S1A-B**), CAP_EIT_ showed comparatively small location-dependent deviations across the three positions, whereas eGFR_EIT_ showed greater within-participant variation (**Fig. 2A-B**). At the middle plane, the median absolute deviation was 0.87% (interquartile range [IQR], 0.62-1.09%) for CAP_EIT_ and 1.23% (IQR, 0.20-7.94%) for eGFR_EIT_ (paired Wilcoxon signed-rank test, p = 0.16) (**Fig. 2C**). At the opposite organ plane, the median deviation remained low for CAP_EIT_ (0.94%; IQR, 0.66-1.70%) but increased significantly for eGFR_EIT_ (2.92%; IQR, 0.46-12.53%; p = 0.008) (**Fig. 2D**). This location sensitivity is consistent with prior EIT studies demonstrating that electrode displacement can alter measured impedance distributions as well as regional information from reconstructed images^34,36^.

**Figure 2.**
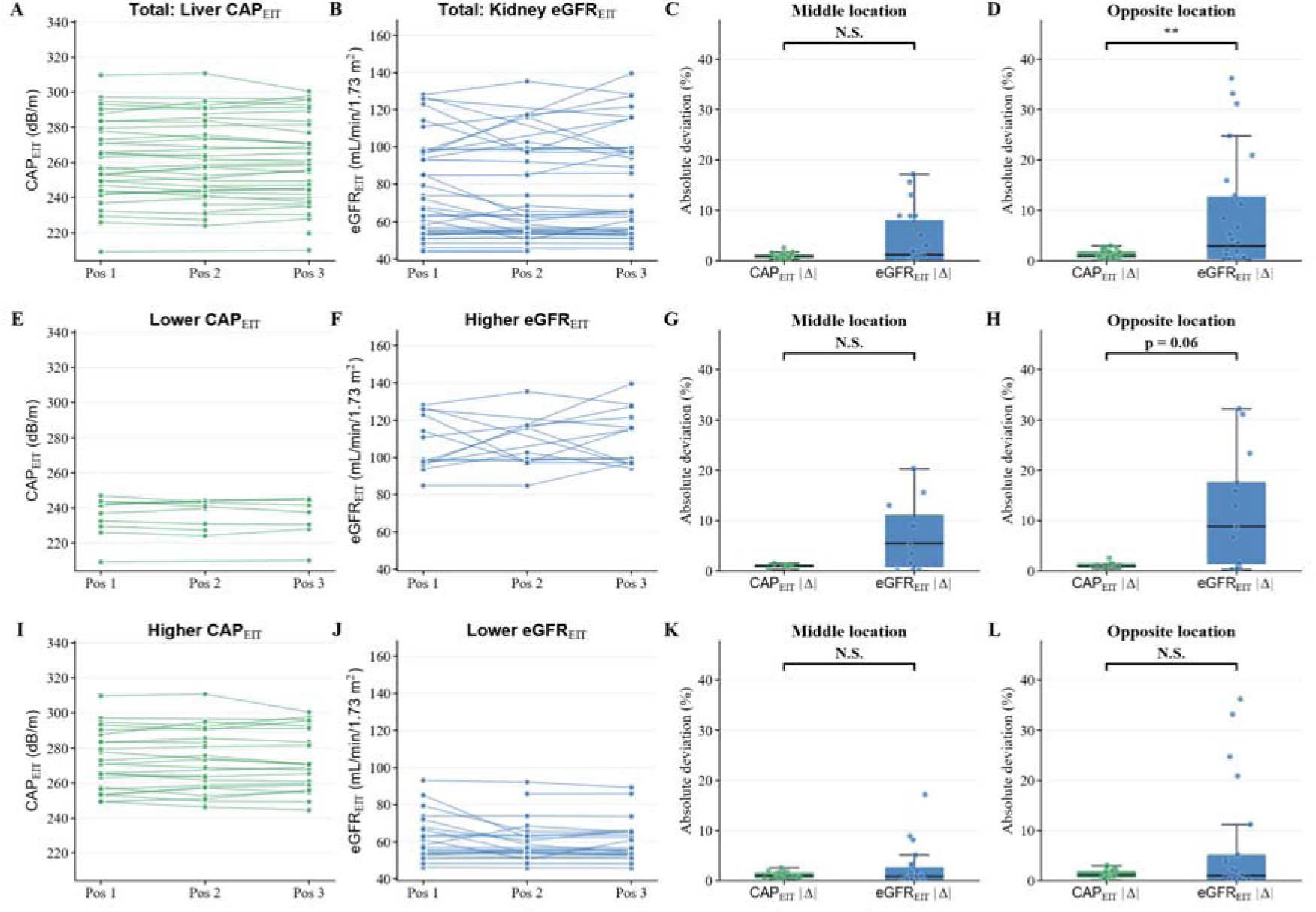
Opposite belt placement produces greater deviation in eGFR_EIT_ than CAP_EIT_. (**A-B**) Participant-level CAP_EIT_ (A) and eGFR_EIT_ (B) estimates at position 1 (Pos 1, conventional liver plane), position 2 (Pos 2), and position 3 (Pos 3, conventional kidney plane) for the full cohort. (**C-D**) Absolute percent deviation from each organ’s conventional belt location at the middle and opposite locations. Two-sided Wilcoxon signed-rank tests, ** p < 0.01. (**E-H**) Corresponding results among participants with lower CAP_EIT_ (≤ 248 dB/m) or higher eGFR_EIT_ (> 90 mL/min/1.73 m^2^) and (**I-L**) among participants with higher CAP_EIT_ (> 248 dB/m) or lower eGFR_EIT_ (≤ 90 mL/min/1.73 m^2^). Mann-Whitney U test, N.S. not significant. Lines in the paired scatter plots connect measurements from the same participant. Boxplots represent the median and interquartile range (IQR), and whiskers extend to the most extreme value within 1.5 times the interquartile range.

Exploratory subgroup analyses classified participants separately according to each organ-health estimate at its conventional acquisition plane. Liver measurements were separated into categories with CAP_EIT_ less than (inclusive) or greater than 248 dB/m using values obtained at position 1. Kidney measurements were separated into categories with eGFR_EIT_ greater than or less than (inclusive) 90 mL/min/1.73 m^2^ using values obtained at position 3. These exploratory cutoffs were selected with reference to commonly reported CAP and eGFR categories but were not used as diagnostic criteria for MASLD or CKD, respectively ^10,37,38^. Within the lower CAP_EIT_ and higher eGFR_EIT_ individuals (**Fig. 2E-F)**, the median absolute percentage deviation at the middle plane was 1.01% (IQR, 0.80 - 1.19%; n = 8) for CAP_EIT_ and 5.43% (IQR, 0.89 - 10.98%; n = 11) for eGFR_EIT_ (Mann-Whitney U, p = 0.11) (**Fig. 2G**). At the opposite plane, the corresponding deviations were 0.89% (IQR, 0.81 - 1.34%; n = 7) and 8.86% (IQR, 1.51 - 17.50%; n = 13), respectively (Mann-Whitney U, p = 0.06) (**Fig. 2H**). Within the higher CAP and lower eGFR groups (**Fig. 2I-J**), median deviations at the middle plane were 0.90% (IQR, 0.60 - 1.44%; n = 21) for CAP_EIT_ and 0.77% (IQR, 0.12 - 2.51%; n = 23) for eGFR_EIT_ (Mann-Whitney U, p = 0.45) (**Fig. 2K**). At the opposite plane, the corresponding deviations were 1.11% (IQR, 0.67 - 1.79%; n = 22) and 0.94% (IQR, 0.27 - 5.00%; n = 22), respectively (Mann-Whitney U, p = 0.55) (**Fig. 2L**). Considering all outputs together, the conventional kidney plane (position 3) provided the most favorable practical compromise where eGFR_EIT_ was measured at its conventional and accurate location, and where CAP_EIT_ remained stable relative to the conventional liver plane.

### Cross-plane image concordance reveals greater kidney sensitivity

We next examined whether the location dependence of the organ-health estimates was accompanied by corresponding changes in reconstructed image structure. Such changes are expected because single-plane EIT technically samples a small, three-dimensional volume that is susceptible to electrode-position and geometry mismatch ^19,39^. For the liver, while raw images showed little variation (**Fig. S2A**), group-averaged difference maps revealed spatially heterogeneous changes across belt locations and frequency pairs (**Fig. 3A**). Despite these differences, images acquired at the middle plane remained more concordant with images from the conventional liver plane than those acquired at the opposite plane. In this regard, middle-plane images exhibited significantly higher spatial correlation (Δz = 0.188, 95% confidence interval [CI], 0.020 - 0.356, p = 0.028) (**Fig. 3B**) and significantly lower normalized root-mean-square error (ΔnRMSE = -0.114, 95% CI, -0.202 to -0.026, p = 0.011) (**Fig. 3C**) than opposite-plane images.

**Figure 3.**
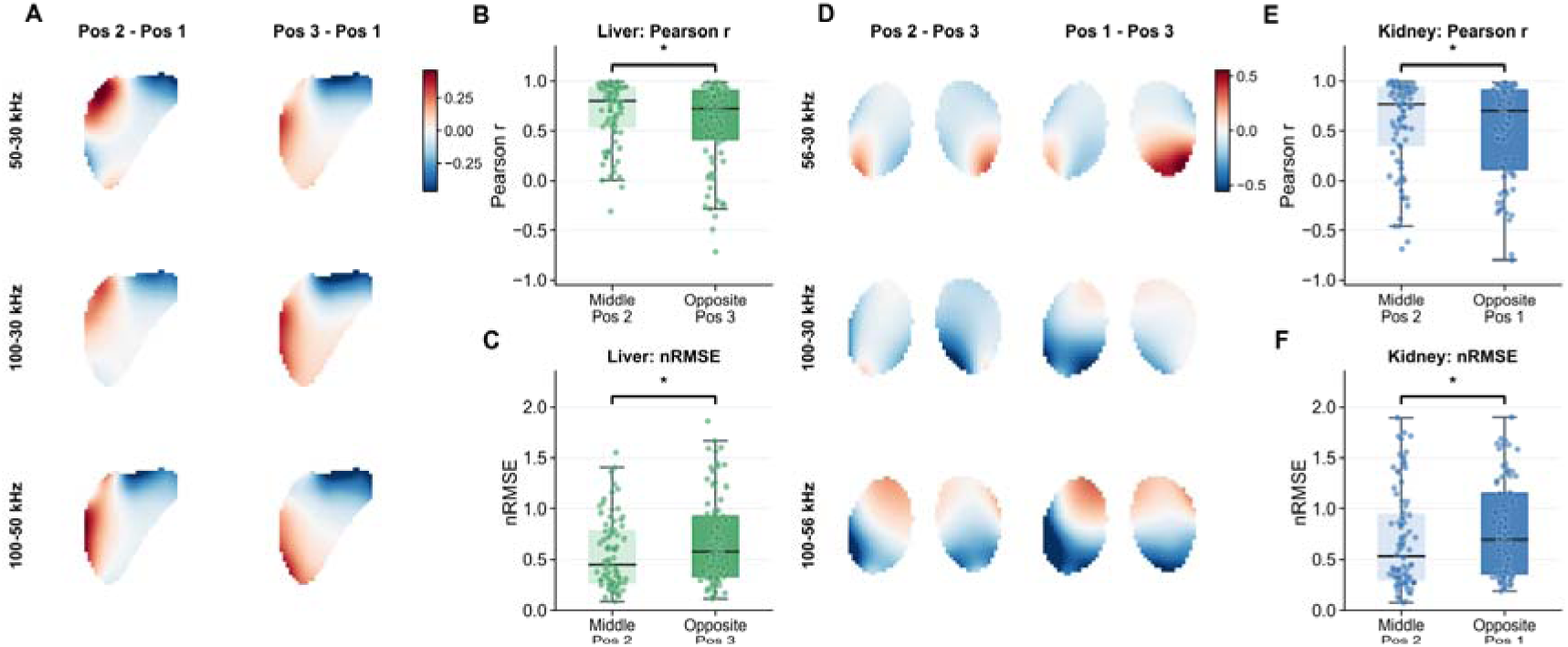
Middle belt placement better preserves organ-specific frequency-difference image structure than opposite placement. **(A)** Group-level differences between liver frequency-difference EIT (fdEIT) images acquired at the middle location (Pos 2) or opposite location (Pos 3) and those acquired at the conventional liver location (Pos 1). Rows show the changes across the 50-30, 100-30, and 100-50 kHz frequency differences. (**B-C**) Spatial Pearson correlation and normalized root-mean-square error (nRMSE), respectively, between liver images acquired at the conventional liver location and those acquired at the middle or opposite location. **(D)** Corresponding kidney image differences between the middle location (Pos 2) or opposite location (Pos 1) and the conventional kidney location (Pos 3). Rows show the changes across the 56-30, 100-30, and 100-56 kHz frequency differences. (**E-F**) Spatial Pearson correlation and nRMSE, respectively, between kidney images acquired at the conventional kidney location and those acquired at the middle or opposite location. Boxplots represent the median and interquartile range (IQR), and whiskers extend to the most extreme value within 1.5 times the interquartile range. Location effects were tested using linear mixed-effects models with belt location and frequency pair as fixed effects and participant as a random intercept. * p < 0.05.

Kidney images similarly exhibited location- and frequency-dependent spatial changes relative to the conventional kidney plane (**Fig. 3D, Fig. S2B**). As observed for the liver, kidney images acquired at the middle plane had significantly higher spatial correlation (Δz = 0.252, 95% CI, 0.047 - 0.458, p = 0.016) (**Fig. 3E**) and lower nRMSE (ΔnRMSE = -0.150, 95% CI, -0.268 to -0.032, p = 0.013) (**Fig. 3F**) than those acquired at the opposite plane. Specifically, the estimated middle-to-opposite contrast was numerically larger for kidney than liver images for both z-scored correlation values (Δz = 0.252 versus 0.188) and the magnitude of the nRMSE difference (|ΔnRMSE| = 0.150 versus 0.114). Although the fitted location contrasts were numerically larger for kidney images, participant-level liver and kidney concordance measures were not significantly different at either the middle or opposite plane (**Fig. S3A-B**).

### Kidney-plane liver estimates remain repeatable after belt repositioning

Considering the two outputs together, position 3 provided the most favorable practical compromise. Liver estimates remained comparatively stable at this plane, whereas kidney estimates were more sensitive to acquisition at the conventional liver plane (position 1). We therefore next tested whether liver estimates from its non-conventional plane remained stable after belt removal and reapplication. At the conventional liver plane, repeated CAP_EIT_ estimates were strongly correlated (r = 0.98, p = 3.4*10^-10^) (**Fig. 4A**), with a mean bias of 1.12 dB/m (paired t-test, p = 0.29) and 95% limits of agreement (LoA) -6.35 to 8.59 dB/m (**Fig. 4B**). These measurements demonstrated that liver health estimates are notably robust when imaging is performed at the liver plane and establish a benchmark for comparison at non-conventional imaging planes. Indeed, repositioning is an important practical stress test because regional measurements derived from EIT images can vary with repeated belt placement in the transverse acquisition plane ^40^.

**Figure 4.**
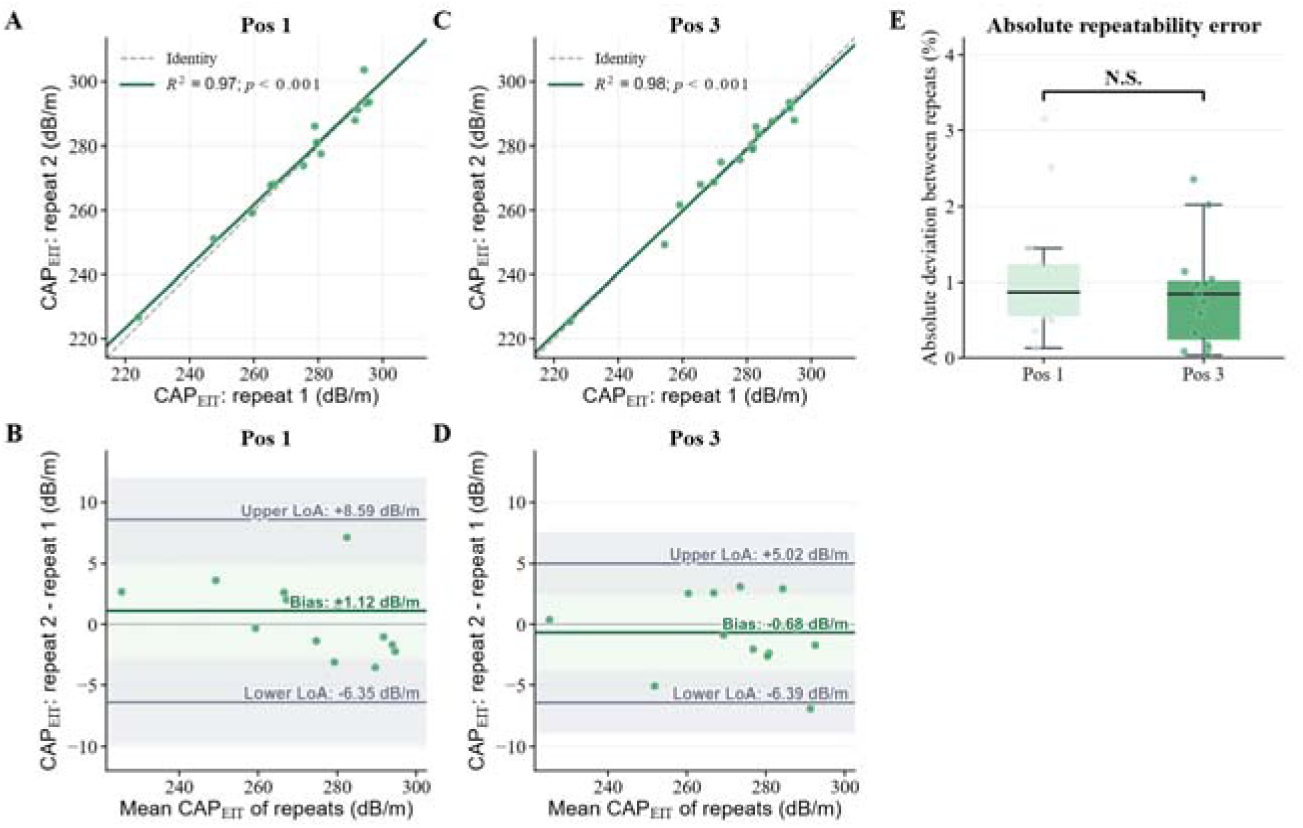
Kidney-plane CAP_EIT_ remains repeatable after belt repositioning. **(A)** Agreement between CAP_EIT_ estimates obtained before and after belt removal and reapplication at Pos 1 (conventional liver position). Points represent individual participants, dashed lines indicate identity, and solid lines indicate linear fits. **(B)** Bland-Altman analyses at Pos 1. Differences are defined as repeat 2 minus repeat 1 and are plotted against the mean of the two measurements. **(C-D)** Corresponding analyses at Pos 3 (conventional kidney position). Green horizontal lines indicate mean bias, dark gray lines indicate the upper and lower 95% limits of agreement calculated as the bias ±1.96 SD, and thin gray lines indicate zero difference. Pale green shading denotes the region between the limits of agreement, whereas gray shaded indicates the approximate 95% confidence intervals around the upper and lower limits of agreement. **(E)** Absolute percentage deviation between repeated CAP_EIT_ estimates at Pos 1 and Pos 3. Boxplots represent the median and interquartile range (IQR), and whiskers extend to the most extreme value within 1.5 times the interquartile range. Two-sided paired Wilcoxon signed-rank comparison, N.S. not significant.

In fact, repeatability at the kidney plane was comparable. Repeated CAP_EIT_ estimates were again highly correlated (r = 0.99, p = 7.9*10^-12^) (**Fig. 4C**), with a mean bias of -0.68 dB/m (paired t-test, p = 0.38) and 95% LoA -6.39 to 5.02 dB/m (**Fig. 4D**). In addition, Bland-Altman proportional-bias checks did not indicate that the repetition difference varied with the magnitude of CAP_EIT_ at either the liver plane (r = -0.13, p = 0.65) or the kidney plane (r = -0.15, p = 0.59). Finally, the absolute percentage deviation between repeats was low and did not differ between positions (Pos 1: median, 0.87%; IQR, 0.57 - 1.22%; Pos 3: median, 0.85%; IQR, 0.25 - 1.01%; paired Wilcoxon signed-rank test, p = 0.67) (**Fig. 4E**), indicating that kidney-plane liver estimates retained the repeatability observed at the liver plane.

### Kidney-plane liver estimates distinguish metabolic risk groups

Having established the robustness and stability of liver health measurements at the kidney plane, we finally examined whether the liver estimates retained clinically relevant information at that location. For this, we leveraged a previous dataset (n = 146) acquired at position 3 that examined the correspondence of eGFR_EIT_ measurements against serum-derived eGFR values ^20,32^. While ground truth CAP values were not measured in this cohort, detailed medical history was acquired that enabled stratification of high and low metabolic liver risk groups. Therefore, this analysis tested the separation of a clinically meaningful metabolic phenotype rather than agreement with a hepatic steatosis diagnosis. To do this, individuals were labelled as high risk if they exhibited both diabetes and a systolic blood pressure ≥140 mmHg (n = 17). By contrast, individuals were labelled as low risk if they exhibited neither (n = 58). 71 of the 146 participants were excluded because they did not meet either definition.

Across the 75 included participants, kidney-plane CAP_EIT_ was significantly higher in the high-risk group (median, 286.0 dB/m; IQR, 281.1 - 303.9 dB/m) than in the low-risk group (median, 267.1 dB/m; IQR, 256.1 - 284.3 dB/m) (two-sided Mann-Whitney U test, p = 1.4*10^-3^) (**Fig. 5A**). In addition, the estimated CAP_EIT_ values discriminated the two groups with a 94.1% sensitivity and 51.7% specificity at the Youden-index cutoff (**Fig. 5B**), and an area under the receiver operating characteristic (ROC) curve (AUC) of 0.756 (95% CI, 0.625 - 0.871) (**Fig. 5C**).

**Figure 5.**
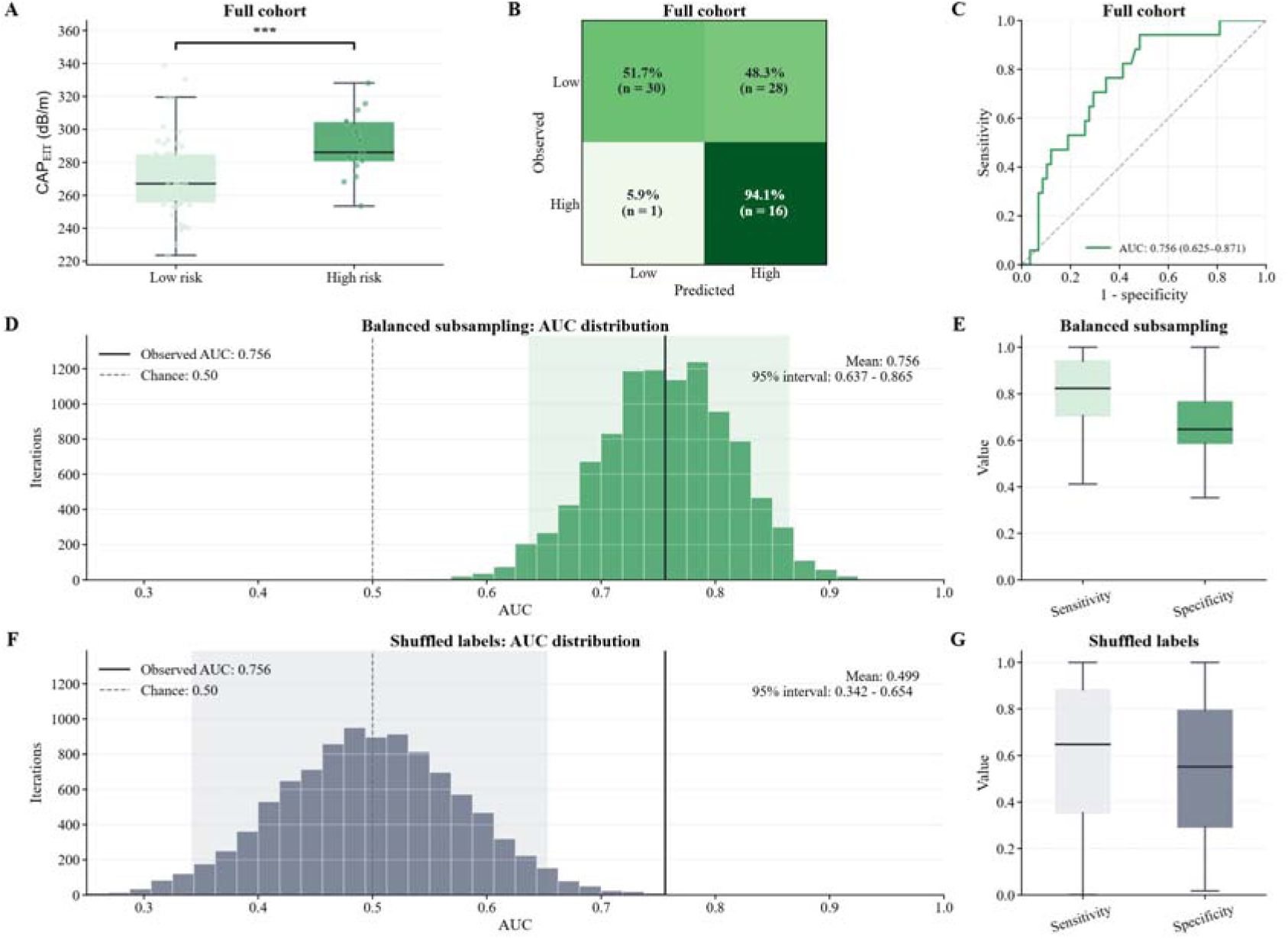
CAP_EIT_ distinguishes high from low metabolic liver-risk groups. **(A)** CAP_EIT_ distributions for high (n = 17) and low (n = 58) metabolic risk groups. High risk was defined by the presence of diabetes together with systolic blood pressure ≥140 mmHg, whereas low risk was defined by the absence of diabetes and systolic blood pressure <140 mmHg. Two-sided Mann-Whitney U tests, *** p < 0.005. **(B)** Confusion matrix for classifying individuals with high vs low metabolic risk. Colors represent percentage. **(C)** Receiver-operating-characteristic curve for the full cohort. The full-cohort AUC confidence interval was estimated using 5,000 bootstrap samples**. (D)** Distribution of AUC values across 10,000 balanced subsamples, each containing 17 high-risk and 17 low-risk participants. **(E)** Distributions of sensitivity and specificity across the balanced subsamples. **(F)** Null distribution of AUC values obtained after permuting the risk labels across 10,000 iterations. **(G)** Distributions of sensitivity and specificity across the label-permutation iterations. In (A), (E) and (G), boxes represent the median and IQR, whiskers extend to the most extreme values within 1.5 times the IQR. In (D) and (F), the solid vertical line indicates the observed full-cohort AUC, the dashed vertical line indicates chance discrimination, and the shaded region denotes the 2.5^th^ - 97.5th percentiles of the subsampling/permutation distribution.

Because the original groups were unbalanced, the analysis was repeated across 10,000 balanced subsamples (n = 17 per group). The mean AUC across these subsamples was 0.756 (95% empirical interval, 0.637 - 0.865) (**Fig. 5D**), and the median sensitivity was 82.4% (IQR, 70.6 - 94.1%) and median specificity 64.7% (IQR, 58.8 - 76.5%; **Fig. 5E**). As a negative control, risk labels were also permuted across 10,000 iterations. The resulting AUC distribution was centered at chance (mean AUC, 0.499; 95% empirical interval, 0.342 - 0.654), and the observed AUC significantly exceeded the distribution of shuffled permutations (p = 7.0*10^-4^) (**Fig. 5F**). Sensitivity and specificity were also broadly distributed after label permutation, with median values falling near chance level, at 64.7% (IQR, 35.3 - 88.2%) and 55.2% (IQR, 29.3 - 79.3%), respectively (**Fig. 5G**). Together, the full-cohort, balanced-subsampling, and permutation analyses indicate that clinically relevant liver-risk information remained detectable from the kidney acquisition plane.

As a sensitivity analysis, we further incorporated self-reported cholesterol status by requiring abnormal cholesterol in the high-risk group and normal cholesterol in the low-risk group. This additional criterion was not included in the primary analysis because cholesterol status was self-reported without clinical confirmation of dyslipidemia and reduced the high- and low-risk groups to 14 and 25 participants, respectively. Nevertheless, CAP_EIT_ remained significantly higher in the high-risk group (Mann-Whitney U test, p = 0.028), with an AUC of 0.716 (95% CI, 0.547 - 0.869) that remained significantly above the label-permutation distribution (p = 0.01) (**Fig. S4A-G**).

## Discussion

In this study, we evaluated whether previously developed liver and kidney EIT pipelines could be deployed from a common abdominal belt plane. We found that the conventional kidney plane provided the most practical compromise for integrated assessment of both organs. CAP_EIT_ remained stable when the belt was moved from the conventional liver plane to the kidney plane, whereas eGFR_EIT_ exhibited significantly greater variation when acquisition moved away from the conventional kidney plane. Reconstructed images from both organs also changed with belt location, with the larger kidney effects consistent with the greater location sensitivity of the downstream eGFR_EIT_ estimate. Liver estimates obtained at the kidney plane also remained highly repeatable after complete belt removal and reapplication, and also retained an association with metabolic liver risk in an independent clinical cohort. Together, these results support the feasibility of extracting complementary liver and kidney information from a common EIT acquisition plane. By removing the need for organ-specific belt repositioning, plane standardization also provides a practical foundation for scalable implementation in primary-care and community-based settings.

A multi-organ approach such as this is highly relevant to clinical workflows because liver and kidney disease commonly arise within the same metabolic context. Contemporary medical frameworks that focus on cardiovascular, kidney, and metabolic disorders emphasize the interaction between metabolic risk factors and organ dysfunction rather than treating each condition as an isolated process ^41^. Meta-analyses and longitudinal cohort studies have also identified a link between steatotic liver disease and subsequent CKD, particularly when metabolic dysfunction is present ^42,43^. With this in mind, the present framework does not aim to replace organ-specific clinical references used to diagnose either condition, but rather addresses an earlier point in the clinical pathway. If one examination can identify complementary liver- and kidney-related risk signals, this information can be used to justify more targeted laboratory or imaging evaluation.

The location sensitivity analysis performed here illustrates an important property of EIT. Although EIT images are commonly displayed as cross-sectional maps, the measured voltages arise from a three-dimensional sensitivity field rather than a cleanly defined anatomical slice ^19^. Consequently, moving the electrode plane changes both the tissues contributing to the measurements and their spatial sensitivity. Furthermore, electrode movement and geometry mismatch are established sources of variation in reconstructed EIT images ^36,39^, and anatomically informed reconstruction can improve the interpretability of regional impedance changes ^44^. Our observation that the middle plane preserved greater image similarity (i.e. high correlation and lower nRMSE) than the opposite plane therefore emphasizes that belt placement remains an important component of abdominal EIT protocol design. The importance of this belt location dependence is also informative in terms of the two organ health estimates. Both liver and kidney images demonstrated significant location-dependent changes, yet CAP_EIT_ remained comparatively stable at the kidney plane while eGFR_EIT_ varied more at the liver plane. One possible explanation is that the larger size of the liver allows better detection of liver-related impedance features across a broader craniocaudal range. Another possibility is that the deep feature embeddings used by the liver analytics pipeline are more robust to electrode displacement than the principal component-based features utilized by the kidney pipelines ^20,31,32^. Regardless, these findings demonstrate that visible image variation does not necessarily produce an equivalent change in a downstream health estimate. Robustness must therefore be evaluated at both the reconstructed-image and model-output levels.

Test-retest repeatability provided a separate test of the proposed kidney-plane implementation. EIT studies in other anatomical settings have shown repeated electrode placement can greatly influence regional measurements ^40^. Despite this well-known source of variability, kidney-plane CAP_EIT_ measurements showed minimal mean bias and narrow limits of agreement after the belt was removed and reapplied. In fact, the repeatability metrics observed here were highly similar to those observed at the conventional liver plane. This is important as clinical CAP measurements themselves can exhibit meaningful measurement variability across operators, examinations and disease stages ^45,46^. While the current CAP_EIT_ repeatability analysis here did not evaluate the ability to repeatedly measure accurate steatosis grades, the high consistency observed is encouraging for establishing the kidney-plane as a robust measurement site.

Finally, the independent clinical analysis provided preliminary evidence that liver-related information remained present at the kidney plane. Specifically, CAP_EIT_ was higher among participants with the combined phenotype of diabetes and elevated systolic blood pressure. These two metabolic features were selected because both are established cardiometabolic risk factors associated with MASLD and more advanced liver disease ^47–49^. For this exploratory analysis, we defined deliberately contrasting metabolic-risk phenotypes with one group of participants classified as high risk when they reported diabetes and had systolic blood pressure ≥140 mmHg, and another group as low risk when they reported no diabetes and had systolic blood pressure <140 mmHg. The observed discrimination even persisted when group size was balanced and disappeared after label shuffling. While dyslipidemia is also an established metabolic risk factor for MASLD ^50^, our clinical data only included self-reported abnormal cholesterol rather than a laboratory-confirmed diagnosis. Nevertheless, when including this additional constraint in a secondary analysis with a smaller high-risk group, the separation between groups persisted. Importantly, this analysis evaluated a metabolic-risk phenotype rather than imaging-confirmed steatosis. The next steps should therefore include an evaluation of the proposed framework prospectively using one standardized kidney-plane acquisition together with clinical CAP or MR-PDFF, as well as serum-derived eGFR measurements for comprehensive multi-organ evaluation. To further integrate liver and kidney imaging into one simultaneous measurement, the frequency and breathing protocols should also be harmonized between the two acquisitions. In this sense, participants are currently instructed to perform a breath hold for liver acquisition and normal breathing for kidney acquisition, however, identifying a single behavior that is suitable for both organs would further optimize screening of both liver and kidney health.

Beyond clinical validation, the proposed single-plane framework creates a practical pathway toward community-scale multi-organ assessment. Eliminating belt repositioning simplifies the acquisition workflow and reduces examination time, operator burden, and opportunities for belt location-related variability. Larger community-based studies should therefore evaluate implementation outcomes alongside liver and kidney risk screening performance, throughput, and the proportion of individuals referred for confirmatory liver or kidney testing. Evaluation across demographically and clinically heterogeneous populations will also be important for determining generalizability. Within such a pathway, EIT would serve as a first-stage risk-assessment tool rather than a diagnostic replacement, allowing portable multi-organ information to guide more targeted use of laboratory testing, elastography, or advanced imaging.

In conclusion, the present results support a portable, single-plane fdEIT framework for integrated liver and kidney assessment. Selecting the kidney plane preserved liver-estimate stability while avoiding the larger displacement-related variation observed for the kidney pipeline. The retained image structure, repositioning repeatability, and independent metabolic-risk association provide complementary evidence that useful liver information remains accessible at this location. With prospective reference standard and external validation, this approach could support scalable multi-organ assessment in primary-care and community settings for identifying individuals who warrant more definitive liver and kidney evaluation.

## Methods

### Study design and participants

The study comprised three analyses of a single-belt strategy for liver and kidney EIT. The belt-location analysis included 42 participants (age range, 18-86 years; 16 male) examined at three adjacent craniocaudal planes. The repeatability analysis included 15 participants (age range, 20-83 years; 3 male) who underwent repeated measurements before and after belt removal. The clinical-risk analysis included 75 participants (age range, 26-93 years; 43 male) from a prior kidney EIT cohort with available metabolic-risk information.

Clinical data were acquired under a protocol approved by the Institutional Review Board of The University of Hong Kong (UW-21-579). Participants provided informed consent before study procedures. Exclusion criteria included recent abdominal surgery, abdominal skin damage, implanted electronic or metallic devices, spinal disease or discomfort, and pregnancy.

### EIT system and acquisition

EIT data were acquired with a portable VitoCheck system (E-SENSE Innovation & Technology Ltd., Hong Kong) using a circumferential 16-electrode belt and a separate reference electrode. A low-amplitude sinusoidal excitation (<1.2 V) was applied sequentially to adjacent electrode pairs, and boundary voltages were recorded from the remaining eligible adjacent pairs. One complete rotation produced 208 measurements per frame (16 current-injection pairs x 13 voltage-measurement pairs). Frames were sampled at 33 Hz.

The liver protocol acquired data at 30, 50 and 100 kHz, and the kidney protocol acquired data at 30, 56 and 100 kHz. Each liver frequency was acquired separately for 7 sec during a breath hold, and each kidney frequency was acquired separately during 12 sec of normal breathing. Three adjacent craniocaudal belt planes were defined relative to the last rib: position 1 above the last rib (the conventional liver plane), position 2 at the last rib (middle plane), and position 3 at the inferior costal margin (the conventional kidney plane). Age, sex, ethnicity, height, weight and waist circumference were also recorded for additional inputs into the pretrained models.

### Faulty electrode detection

Raw voltage data were screened using an established faulty-electrode classifier trained with clean acquisitions and acquisitions in which individual electrodes were intentionally detached ^20^. Measurement-derived features were classified at the frame and frequency level, and frames containing any detected faulty electrode were excluded. Retained frames were averaged within frequency to obtain one measurement vector per frequency. Recordings in which no frequencies survived were excluded from further analysis.

### Organ-health estimates

CAP_EIT_ and eGFR_EIT_ were generated with pretrained models described in the earlier liver and kidney validation studies ^20,31,32^. Reconstructed fdEIT images were converted into model features. The liver pipeline used learned image embeddings and pixel-level features, and the kidney pipeline used principal-component features. EIT features were combined with prespecified demographic and anthropometric variables and entered into pretrained ensemble regressors. Outputs were expressed in the clinical units of the corresponding reference measurements: dB/m for CAP and mL/min/1.73 m^2^ for eGFR. The analysis applied upper output plausibility filters of 400 dB/m for CAP_EIT_ and 150 mL/min/1.73 m² for eGFR_EIT_. The CAP threshold corresponds to the established 100 - 400 dB/m range of clinical CAP measurements, whereas the eGFR threshold was selected at the upper end of the range containing most published definitions of renal hyperfiltration ^51,52^.

### Belt-location sensitivity analysis

Each participant underwent both liver and kidney acquisition protocols at three craniocaudal belt positions, yielding six intended recordings per participant. CAP_EIT_ and eGFR_EIT_ values were calculated independently at each position using the corresponding pretrained organ-health pipelines. Position 1 (the conventional liver position) served as the within-participant reference for CAP_EIT_ while position 3 (the conventional kidney position) served as the reference for eGFR_EIT_. For each non-conventional position, the absolute percentage change was calculated relative to the corresponding conventional position. Deviations were evaluated at the “middle” position and at the “opposite” organ position, defined as position 3 for CAP_EIT_ and position 1 for eGFR_EIT_. Within-participant differences between CAP_EIT_ and eGFR_EIT_ absolute percentage deviations were evaluated separately at the middle and opposite positions using two-sided Wilcoxon signed-rank tests.

Exploratory subgroup analyses also examined whether belt-position sensitivity differed according to organ-health estimates obtained at their conventional positions. Liver measurements were classified as lower CAP_EIT_ (≤248 dB/m) or higher CAP_EIT_ (>248 dB/m) using values obtained at position 1. Similarly, kidney measurements were classified as higher eGFR_EIT_ (>90 mL/min/1.73 m^2^) or lower eGFR_EIT_ (≤90 mL/min/1.73 m^2^) using values obtained at position 3. For the lower-risk comparison, CAP_EIT_ deviations among participants with lower CAP_EIT_ were compared with eGFR_EIT_ deviations among participants with higher eGFR_EIT_. For the higher-risk comparison, CAP_EIT_ deviations among participants with higher CAP_EIT_ were compared with eGFR_EIT_ deviations among participants with lower eGFR_EIT_. Comparisons were performed using two-sided Mann-Whitney U tests.

### Frequency-difference EIT image reconstruction and analysis

Frequency-difference EIT images were generated for all pairwise combinations of the acquired frequencies, with the lower-frequency voltage profile serving as the reference. Liver images were therefore reconstructed for the 50-30, 100-30 and 100-50 kHz frequency pairs, whereas kidney images were reconstructed for the 56-30, 100-30 and 100-56 kHz frequency pairs. Conductivity changes were reconstructed using a regularized Jacobian solver with generic organ-specific meshes. Element-wise reconstructions were projected onto a 64 x 64 rectangular grid, and subsequent analyses were restricted to organ-specific regions of interest derived from the corresponding meshes.

For each participant and frequency pair, images acquired at the two non-conventional positions were compared with the image acquired at that organ’s conventional position. Position 1 served as the reference for liver images, whereas position 3 served as the reference for kidney images. Spatial concordance was quantified using the Pearson correlation coefficient and normalized root-mean-square error (nRMSE). nRMSE was calculated after independently dividing each image vector by its root-mean-square amplitude within the organ-specific region of interest. Pearson correlation coefficients were Fisher-z transformed. Fisher-z correlation and nRMSE were analyzed separately for each organ using linear mixed-effects models containing belt position and frequency pair as fixed effects and participant as a random intercept. Models were fitted using restricted maximum likelihood. Fixed-effect coefficients are reported with two-sided Wald p values and 95% Wald confidence intervals.

For between-organ comparisons, concordance measures were averaged across all three frequency pairs within each participant, organ, and location. Liver and kidney measures were compared within matched participants separately at the middle and opposite planes using two-sided Wilcoxon signed-rank tests, with Holm correction across the two locations and within each metric.

### Test-retest repeatability

Participants in the repeatability analysis underwent two liver acquisitions at both position 1 (the conventional liver plane) and position 3 (the conventional kidney plane), with belt removal and repositioning between repetitions. CAP_EIT_ was calculated independently for each acquisition. Repeatability was summarized at each location using Pearson correlation and Bland-Altman analysis between the first and second repetition ^53^. Mean bias and 95% limits of agreement were calculated as the mean difference ±1.96*SD (standard deviation) of the paired differences. Approximate 95% confidence intervals for each limit of agreement were calculated as the estimated limit 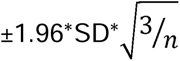 where SD denotes the standard deviation of the paired differences and *n* the number of paired samples. Fixed bias was tested using a two-sided one-sample t-test of the paired differences against zero. Proportional bias was assessed using the Pearson correlation between the participant-level mean and difference. Absolute repeatability error was calculated as the absolute difference between repetitions divided by their mean and expressed as a percentage. Errors at positions 1 and 3 were compared using a two-sided Wilcoxon signed-rank test.

### Metabolic-risk phenotype analysis

Exploratory clinical relevance of kidney-plane liver measurements was examined using a previous clinical cohort of 146 individuals examining EIT-based kidney health^20^. No ground truth CAP values were collected from these individuals, and therefore the analysis evaluated the separation of a predefined metabolic liver-risk phenotype rather than agreement with a diagnosis of hepatic steatosis. Diabetes status was obtained from the clinical questionnaire, and systolic blood pressure was measured during the study visit. Diabetes and elevated blood pressure were selected because both are established metabolic factors associated with steatotic liver disease and more advanced liver disease ^54–56^. A systolic threshold of 140 mmHg was selected to be consistent with that used in the 2024 ESC hypertension definition ^57^. For this exploratory analysis, participants were therefore classified as high risk when they reported diabetes and had a systolic blood pressure ≥140 mmHg, and as low risk when they did not report diabetes and had a systolic blood pressure <140 mmHg. Participants with discordant diabetes and blood-pressure profiles were excluded from this analysis. The resulting analysis cohort comprised 17 high-risk and 58 low-risk participants, with 71 not meeting either criterion.

CAP_EIT_ distributions were compared between the two independent groups using a two-sided Mann-Whitney U test. Discrimination was quantified using the area under the receiver-operating-characteristic curve (AUC). A 95% percentile bootstrap confidence interval for the full-cohort AUC was calculated from 5,000 participant-level bootstrap samples. For descriptive classification metrics, the CAP_EIT_ threshold maximizing Youden’s J statistic was identified from the full cohort and used to calculate sensitivity and specificity.

To evaluate the influence of group imbalance, the analysis was repeated across 10,000 balanced subsamples. For each iteration, 17 low-risk and 17 high-risk participants were sampled without replacement, and the AUC, Youden threshold and sensitivity and specificity were recalculated. The AUC distribution was summarized using its mean and 2.5^th^ - 97.5th percentiles, whereas the sensitivity and specificity distributions were summarized using their medians and IQRs.

As a negative control, the risk labels of the full cohort were randomly permuted 10,000 times while retaining the observed CAP_EIT_ values. The AUC, Youden threshold and sensitivity and specificity were recalculated within each permutation. The null AUC distribution was summarized using its mean and 2.5th - 97.5th percentiles, whereas sensitivity and specificity were summarized using their medians and IQRs. The permutation p value was calculated as

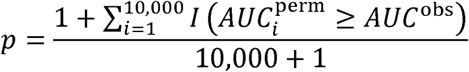

A fixed random seed of 42 was used for all bootstrap, subsampling and permutation procedures.

The same analyses were repeated using a more restrictive three-factor phenotype. In this case, high risk required diabetes, systolic blood pressure ≥140 mmHg, and self-reported abnormal cholesterol, whereas low risk required the absence of diabetes, self-reported normal cholesterol, and systolic blood pressure <140 mmHg. This sensitivity analysis included 14 high-risk and 25 low-risk participants, with 14 participants per group in balanced subsampling.

### Statistical analysis

Continuous variables are summarized as the median and IQR unless otherwise stated. All statistical tests were two-sided, with p < 0.05 considered statistically significant. Analysis-specific statistical tests, models, confidence intervals and resampling procedures are described in the corresponding Methods sections. P values and 95% confidence or empirical intervals are reported where applicable.

## Data and code availability

De-identified derived data sufficient to reproduce the primary figures and statistical analyses, together with analysis code, will be deposited in a public repository upon publication.

## Acknowledgements

This work was supported in part by the Mitzi, Lee Leung Yin Ping Charitable Foundation and E-SENSE Innovation & Technology internal funds. The authors would like to thank all members of E-SENSE Innovation & Technology for helpful discussions related to this work.

## Author Contributions

Conceptualization: BJE, RWC; Data Curation: AW, JCL, MKH, JQJG, BJE; Formal Analysis: AW, JCL, MKH, BJE; Funding Acquisition: MW, RWC; Investigation: AW, JCL, MKH, JQJG; Methodology: NJG, IYZ, BJE, RWC; Project Administration: BJE, RWC; Resources: MW, BJE, RWC; Software: AW, JCL, MKH, JQJG, BJE; Supervision: IYZ, BJE, RWC; Validation: AW, JCL, MKH, BJE; Visualization: AW, JCL, MKH, BJE; Writing - Original Draft: AW, BJE, RWC; Writing - Review & Editing: AW, JCL, MKH, JQJG, NJG, IYZ, MW, BJE, RWC

