## Supplementary materials for "A single-plane electrical impedance tomography framework for integrated liver and kidney assessment"

**This file contains:**

Supplementary Figures 1 - 4

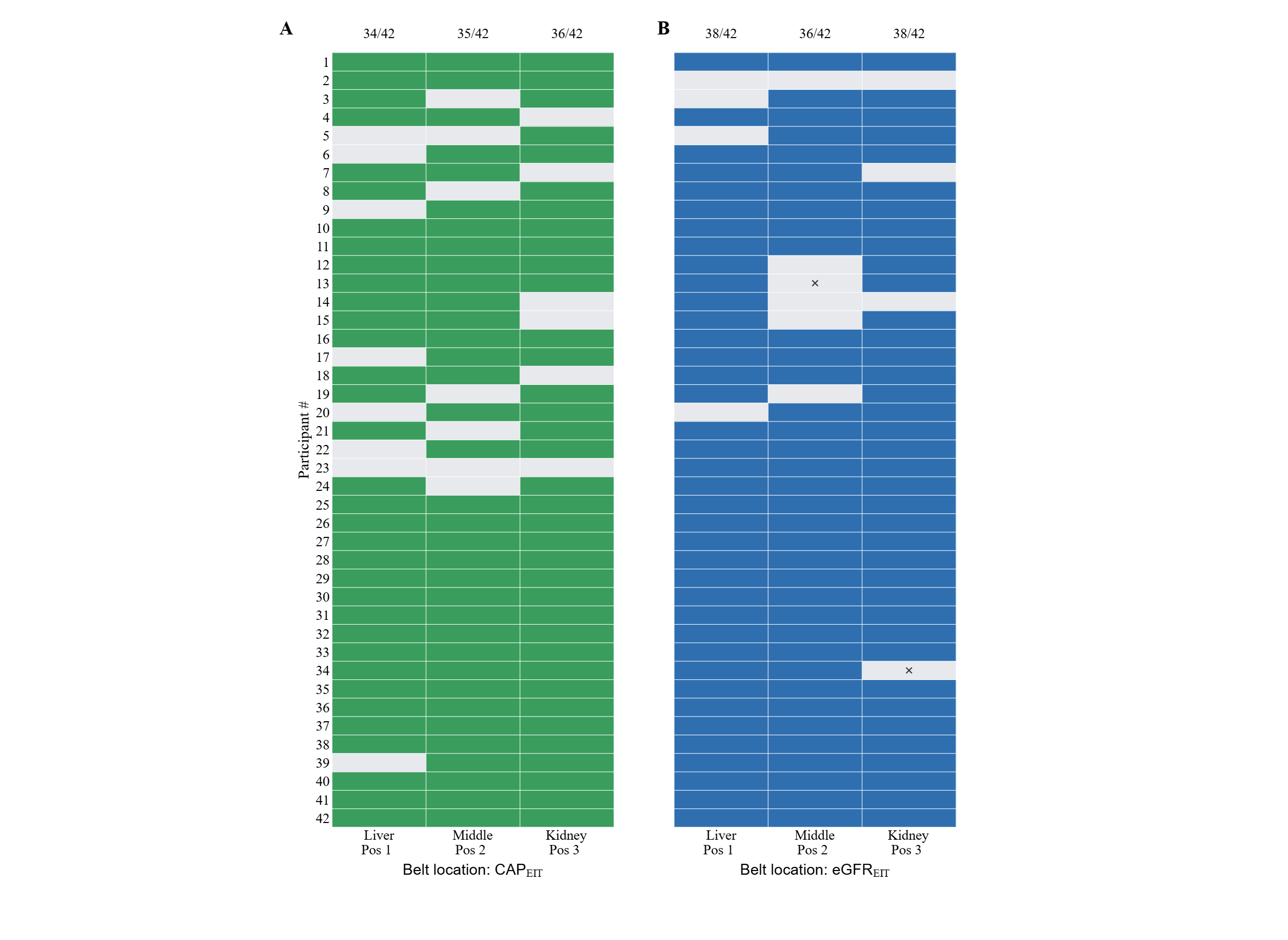

**Figure S1. Availability of organ-health estimates across belt positions.** Valid organ health estimates at the three belt locations for **(A)** CAP_EIT_ and **(B)** eGFR_EIT._ Rows represent participants and columns represent belt positions. Gray cells indicate measurements that did not pass the EIT quality control, and for which no organ-health estimate was computed. Crosses identify prediction values that were excluded because they exceeded the upper output plausibility filters for the respective organ-health metric. Counts above each column indicate the number retained out of 42 participants.

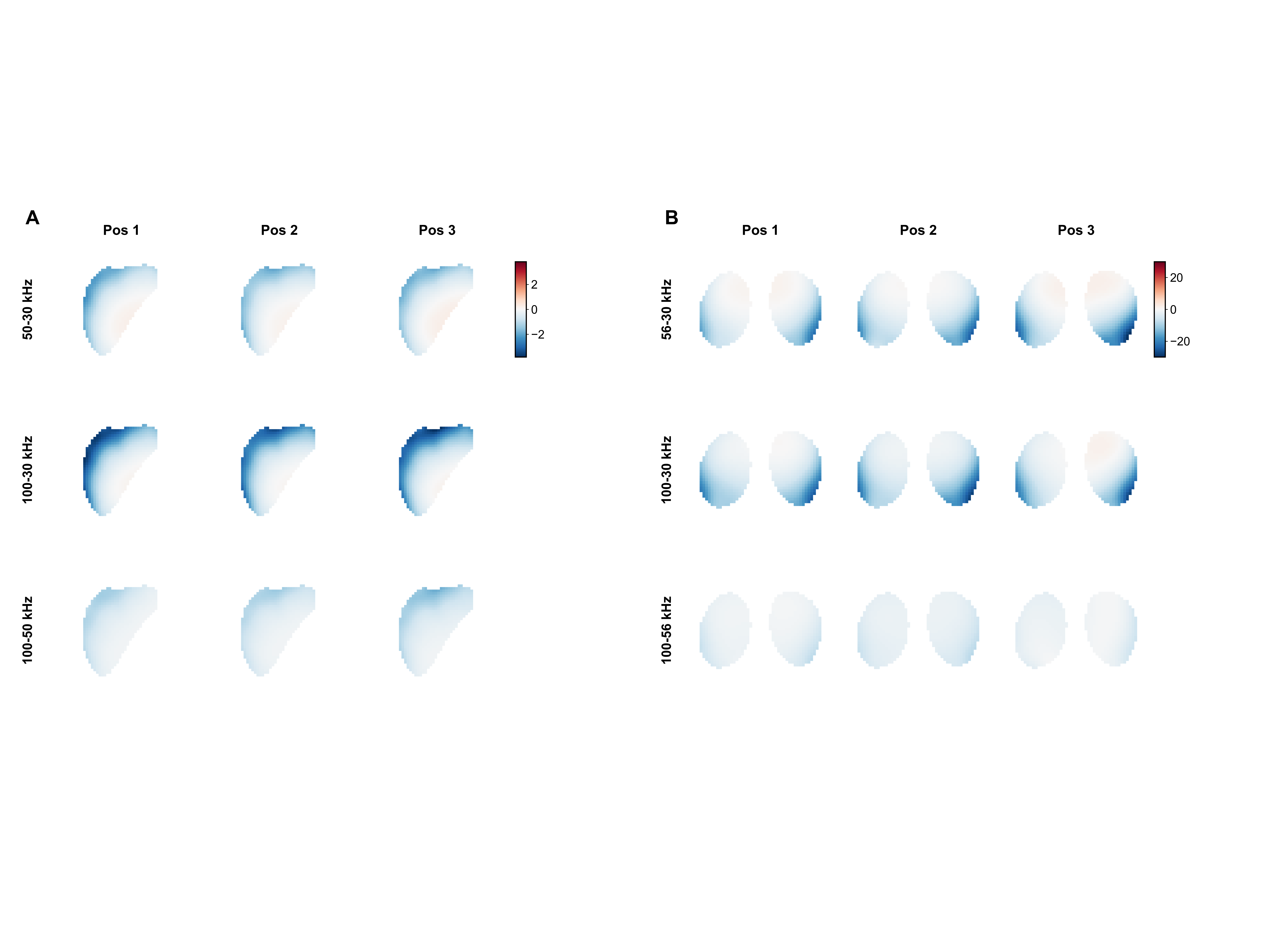

**Figure S2. Reconstructed liver and kidney fdEIT images across belt locations. (A)** Group-averaged liver frequency-difference EIT images at positions 1 (liver), 2 (middle), and 3 (kidney) for the 50-30, 100-30 and 100-50 kHz frequency pairs. **(B)** Corresponding kidney images for the 56-30, 100-30 and 100-56 kHz frequency pairs.

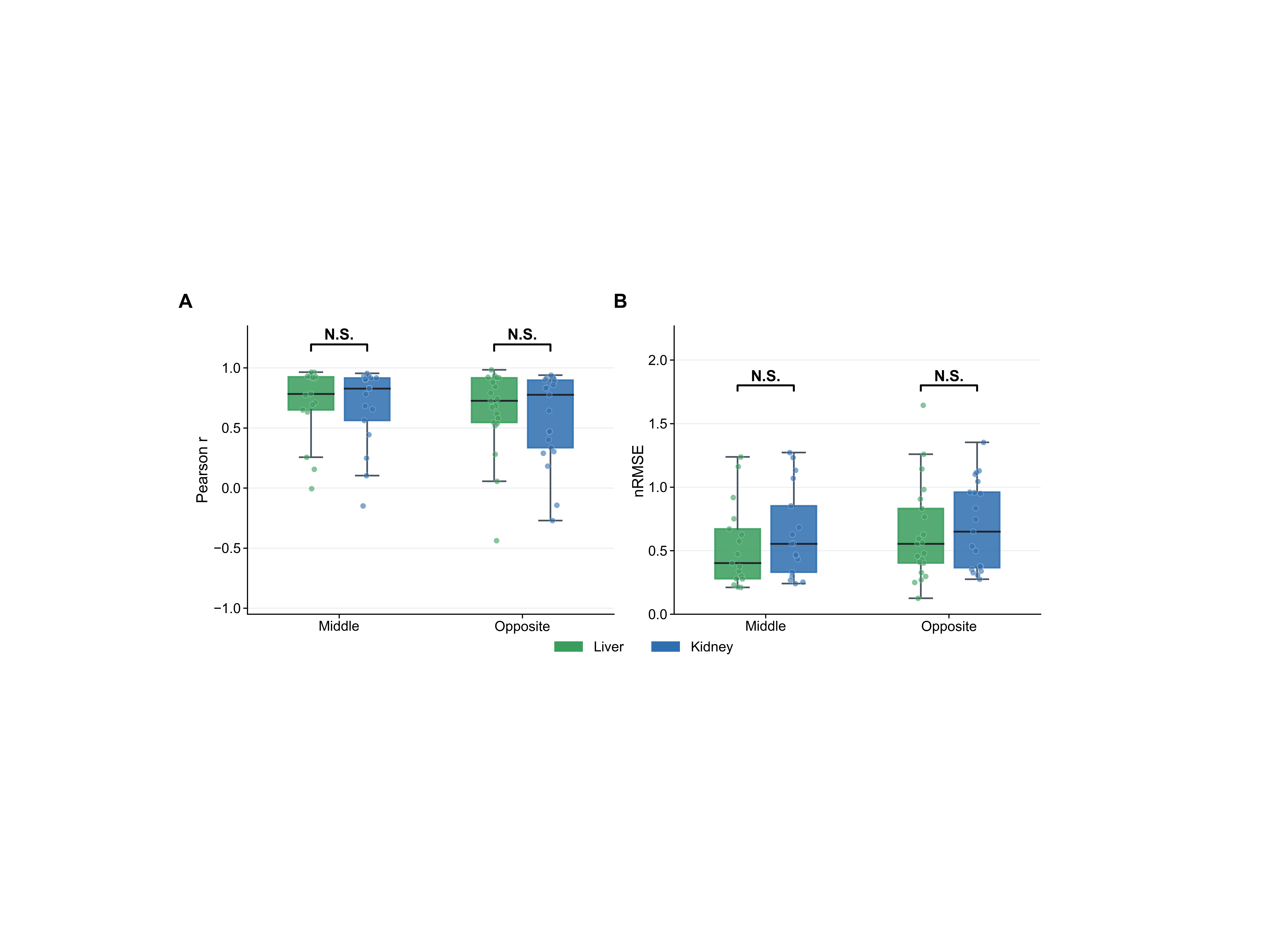

**Figure S3. No significant between-organ differences in image concordance at the middle or opposite planes. (A)** Spatial Pearson correlation and **(B)** normalized root-mean-square error (nRMSE) relative to each organ’s conventional-plane image. Green and blue indicate liver and kidney measurements, respectively. Boxplots represent the median and interquartile range (IQR), and whiskers extend to the most extreme value within 1.5 times the interquartile range. Between-organ comparisons used two-sided Wilcoxon signed-rank tests with Holm correction across locations within each metric. N.S., not significant.

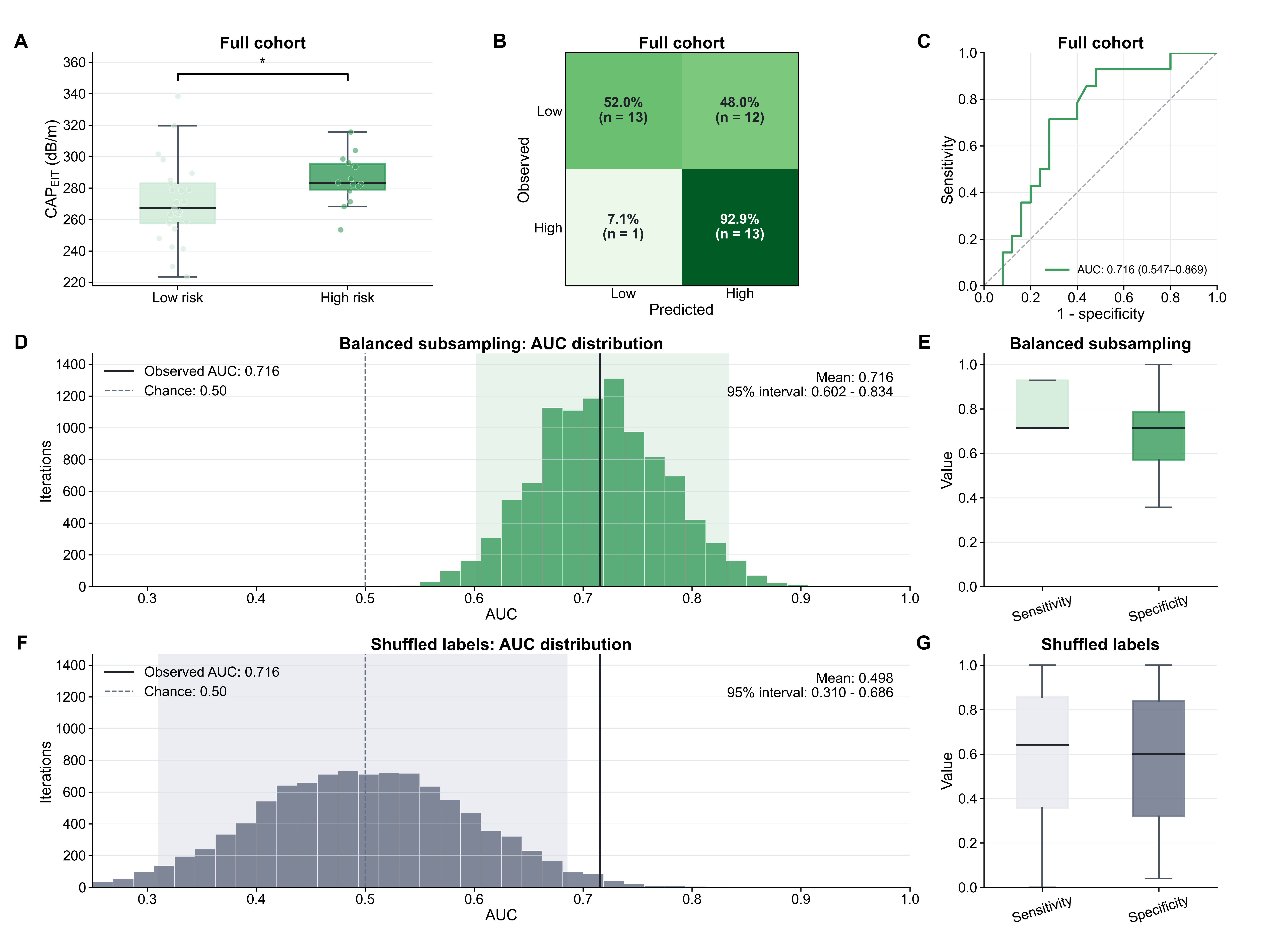

**Figure S4. CAP_EIT_ distinguishes high from low metabolic liver-risk groups when self-reported cholesterol status is also required. (A)** CAP_EIT_ distributions for high (n = 14) and low (n = 25) metabolic risk groups. High risk was defined by the presence of diabetes together with systolic blood pressure ≥140 mmHg and self-reported abnormal cholesterol levels, whereas low risk was defined by the absence of diabetes, normal self-reported cholesterol levels, and systolic blood pressure <140 mmHg. Two-sided Mann-Whitney U tests, * p < 0.05. **(B)** Confusion matrix for classifying individuals with high vs low metabolic risk. Colors represent percentage. **(C)** Receiver-operating-characteristic curve for the full cohort. The full-cohort AUC confidence interval was estimated using 5,000 bootstrap samples**. (D)** Distribution of AUC values across 10,000 balanced subsamples, each containing 14 high-risk and 14 low-risk participants. **(E)** Distributions of sensitivity and specificity across the balanced subsamples. **(F)** Null distribution of AUC values obtained after permuting the risk labels across 10,000 iterations. **(G)** Distributions of sensitivity and specificity across the label-permutation iterations. In (A), (E) and (G), boxes represent the median and IQR, whiskers extend to the most extreme values within 1.5 times the IQR. In (D) and (F), the solid vertical line indicates the observed full-cohort AUC, the dashed vertical line indicates chance discrimination, and the shaded region denotes the 2.5^th^ - 97.5th percentiles of the subsampling/permutation distribution.
